# Exploring serial crystallography for drug discovery

**DOI:** 10.1101/2023.12.21.572778

**Authors:** A. Dunge, C. Phan, O Uwangue, M. Bjelcic, J. Gunnarsson, G. Wehlander, H. Käck, G. Brändén

## Abstract

Structure-based drug design is highly dependent on the availability of structures of the protein of interest in complex with lead compounds. Ideally, this information can be used to guide the chemical optimization of a compound into a pharmaceutical drug candidate. A limitation of the main structural method used today, rotational cryo-crystallography, is that it only provides structural information of the protein-complex in its frozen state. Serial crystallography is a relatively new approach that offers the possibility to study protein structures at room-temperature. Here, we explore the use of serial crystallography to determine structures of the pharmaceutical target soluble epoxide hydrolase. We introduce a new method to screen for optimal microcrystallization conditions suitable for use in serial crystallography and present a number of room-temperature ligand-bound structures of our target protein. From a comparison between the room-temperature structural data and previously published cryo-temperature structures, we describe an example of a temperature-dependent difference in ligand-binding mode and observe that flexible loops are better resolved at ambient temperature. Finally, we discuss current limitations and potential future advances of serial crystallography for use within pharmaceutical drug discovery.

## 1. Introduction

Structure-based drug design is a very well-established concept, integral to drug discovery since it improves the speed and quality of pharmaceutical candidate compound discovery. Structural methods are commonly used throughout a drug-design project from early hit-finding to optimization of potency and properties for *in-vivo* delivery (Maveyraud & Mourey, 2020). Thus, all major pharmaceutical companies and many biotech companies have access to structural biology capabilities. Successful structure-based drug design requires structure determination to high quality on a timescale matching that of the design cycle. Conventional macromolecular X-ray crystallography, performed on single protein crystals kept at cryogenic temperatures, is highly streamlined and the work horse of structure-based drug design. It does, however, require a very robust crystallization protocol and includes the labor intense step of harvesting crystals. Single-particle cryo-electron microscopy (cryo-EM) has gained much attention during the last years as an attractive alternative to X-ray crystallography as it does not require that the protein of interest is prone to form well-ordered crystals (Guaita *et al*., 2022). However, both traditional cryo-crystallography and single-particle cryo-EM are limited in that they only provide structural information of the target protein in a frozen state.

Serial crystallography (SX) is a relatively new concept where X-ray diffraction data are collected on micrometer-sized crystals at room-temperature. The method was first developed for use at X-ray free electron laser facilities, where the extreme intensity of the X-ray pulses required a new way of collecting diffraction data. To circumvent this problem, injection devices were developed that supply a stream of very small crystals across the X-ray beam such that each crystal that intercepts the X-rays gives rise to one diffraction image (Chapman *et al*., 2011). Pioneering experiments established that although the data recorded is built up of partial reflections from randomly oriented crystals, it is possible to obtain high quality structural information by the use of serial crystallography (Boutet *et al*., 2012). Since then, the concept has been further developed at synchrotron facilities (Gati *et al*., 2014), giving rise to the term serial synchrotron crystallography. There are now a number of purpose-built synchrotron beamlines which allow many more users to explore the method than what the XFEL facilities can cater for. The growth of the method has also led to a wide variety of sample delivery methods being developed. This includes different variants of jets, including the high-viscosity extruder injector that is suitable for viscous samples such as membrane proteins crystallized in lipidic cubic phase (Weierstall *et al*., 2014). There is also a multitude of different fixed-target devices available where the crystals are dispensed onto a surface that is translated across the X-ray beam in a grid-like fashion. This includes highly specialized chips where the crystal surrounding atmosphere can be controlled (Roedig *et al*., 2017), as well as simple X-ray transparent membranes that are sandwiched around the crystals and mounted onto a standard crystallography pin (Roedig *et al*., 2017, 2016; Owen *et al*., 2017), enabling data collection at any high-focus synchrotron beamline equipped with a fast translating goniometer. A large advantage of the fixed target devices is that sample consumption can typically be minimized compared to the jets, a prerequisite for SX to be a realistic alternative for the majority of proteins of interest.

A scientific area where SX has had great impact is within time-resolved structural studies, where the method has opened up unique possibilities previously out of reach by traditional cryo-crystallography (Brändén & Neutze, 2021). Of more general interest, it has been suggested that it may be important to study protein structures at room temperature as opposed to in their frozen state to reduce the risk of temperature artifacts. Several studies have made interesting observations of structural differences related to temperature (Fenwick *et al*., 2014; Ebrahim *et al*., 2022; Keedy *et al*., 2018), including the loss of information about transient binding sites when working at cryogenic temperature (Fischer *et al*., 2015). Very recently, a crystallographic screen of 143 small-compound fragment binders targeting protein tyrosine phosphatase PTP1B was performed using rotational crystallography at room temperature (Skaist Mehlman *et al*., 2023). By comparison with earlier cryo-crystallography data, it was shown that fewer ligands bound at room-temperature, but also that unique binding-poses were identified in the room-temperature experiment including interactions at new binding sites that had previously not been observed. Finally, SX has a potential advantage in that it may be possible to avoid the manual handling of crystals that is required in traditional cryo-crystallography where the protein crystals are typically harvested and manipulated by hand. A future automation of the crystal handling would be highly interesting, not the least for use in high-throughput drug-discovery campaigns.

To explore the utility of SX in a drug-discovery setting, we selected a pharmaceutically relevant target, soluble Epoxide Hydrolase (sEH) as a model system. sEH is an enzyme that binds epoxides and converts them to their corresponding diols (Newman *et al*., 2003). Three residues within the active site, ASP335, TYR383 and TYR466, make up the catalytic triad (Figure 1). The inhibition of sEH can effectively maintain endogenous epoxyeicosatrienoic acids (EETs) levels and reduce dihydroxyeicosatrienoic acids (DHETs) levels, thus providing a therapeutic potential for cardiovascular, central nervous system, and metabolic diseases (Imig & Hammock, 2009; Chiamvimonvat *et al*., 2007; Sun *et al*., 2022; Shen & Hammock, 2012; Hashimoto, 2019; Vázquez *et al*., 2023). Structures of sEH in complex with a large number of inhibitors have been solved previously using conventional cryo-crystallography (Öster *et al*., 2015). Thus, we set out to generate a set of room-temperature protein-inhibitor complexes to compare with those previously solved and explore whether the use of SX would provide additional structural information of interest in a drug-discovery setting.

**Figure 1.**
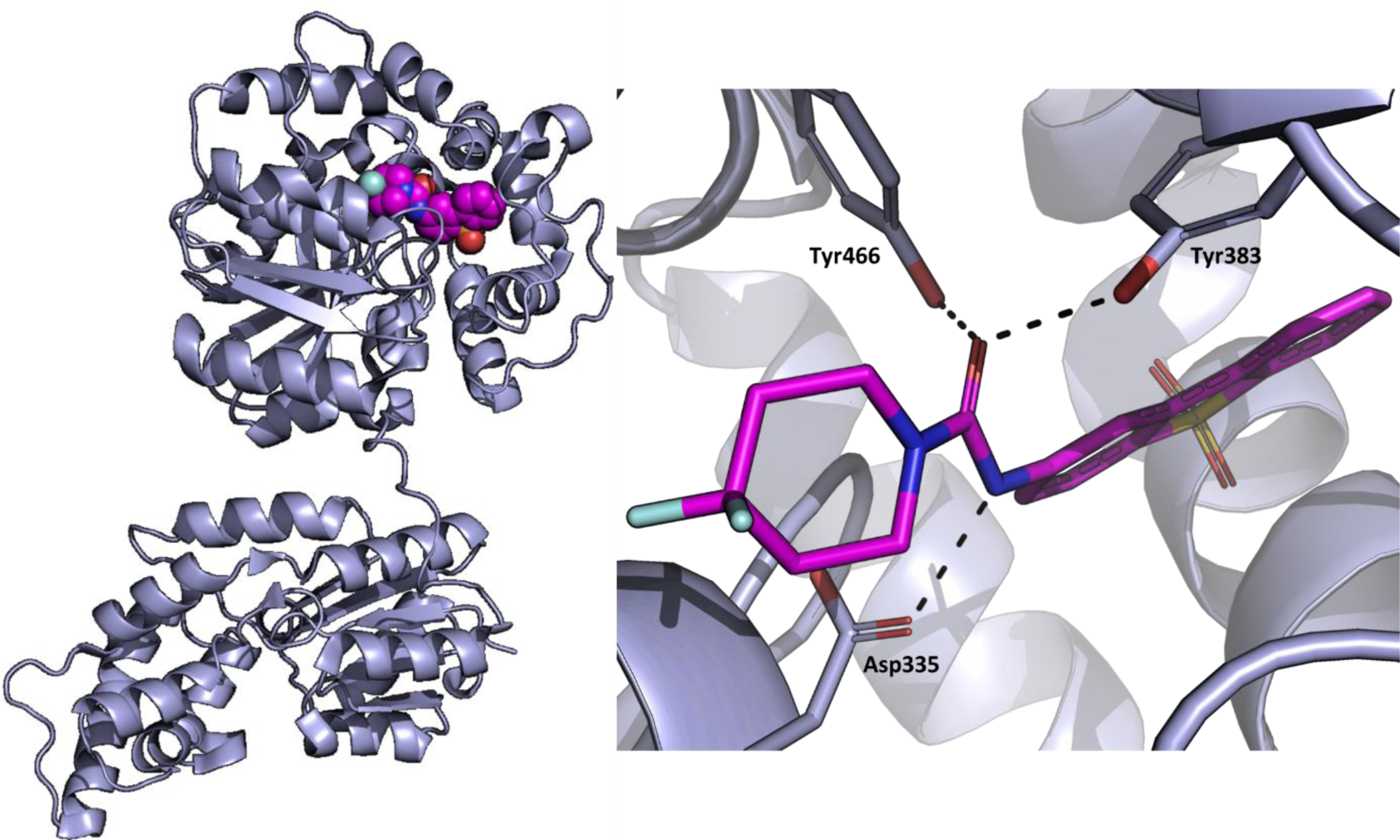
Structure of sEH. The left-hand side panel shows a structure of inhibitor-bound sEH solved at cryogenic temperature (PDB ID 5AKE). The active site with the bound compound (purple) is located in the C-terminal domain. The right-hand side panel shows a zoom-in of the active site where the compound (purple) interacts with the catalytic triad (Asp335, Tyr383 and Tyr466).

## 2. Methods

### 2.1. Protein production and purification

A C-terminally truncated construct of sEH(1-548) was expressed in *Spodoptera frugiperda* (*SF9*) insect cells as previously described (Öster *et al*., 2015). 100 ml of cells were used for each purification following the published protocol with the exception that two interconnected 5 ml HisTrap FF Crude (Cytivia) columns were used instead of one. The purified protein was concentrated to 15-20 mg/ml and stored in a buffer containing 20 mM Tris-HCl pH 8, 150 mM NaCl, 1 mM TCEP and 10 % glycerol.

### 2.2. Crystallization

A concentrated seed solution was generated by growing larger crystals of sEH, 100-1000 µm in size, following a previously established protocol (Öster *et al*., 2015). Briefly, vapor diffusion crystallizations were set up using a protein concentration of 15-20 mg/ml and a 1:1 ratio of protein:precipitant solution in 24-well sitting drop plates (Cryschem M Plate, Hampton research) with a well solution containing 32-38 % PEG3350, 0.1 M LiSO_4_, 0.1 M Tris pH 8.5 and a drop volume of 5-10 µl. The drops were streak-seeded using crystal seeds previously obtained using this protocol. Drops containing well-formed crystals were transferred into a 2 ml Eppendorf tube, typically resulting in a volume of ∼100 µl. Two microseed beads (Molecular Dimensions) were added to the tube and crystals were crushed by vortexing the sample for 3 x 5 min, or until no large fragments of crystals were visible in an optical microscope. This seed production protocol was inspired by a previously published method to produce seeds for microcrystallization (Dods *et al*., 2017).

Optimization of batch crystallization conditions were done in a micro-batch setting using sitting drop plates, referred to as hybrid crystallization. This was achieved by composing the well solution such that the concentration of the individual components corresponds to the concentration in the crystallization drop after addition of protein. A crystallization screen was carried out in this fashion by varying the seed concentration and seed volume. In addition, the concentration of protein, PEG3350, Li_2_SO_4_ and Tris-HCl (pH 8.5) were optimized, as well as the ratio of protein to precipitant solution (see Table S1 for details).

Optimized microcrystals for serial crystallography data collection were produced as follows (Figure 2). A crystallization solution was prepared by mixing 10 % (v/v) seed solution into a precipitant solution containing 34 % PEG3350, 0.1 M Tris-HCl (pH 8.5) and 0.1 M Li_2_SO_4_. Protein at 14 mg/ml was then mixed with the crystallization solution at a 1:4 protein:precipitant/seed solution ratio in a 0.5 ml Eppendorf tube and the solution was vortexed. The total microcrystallization volume was typically 50-100 µl. The sample was kept upright to allow crystals to sediment to the bottom of the tube and stored at 20 ^◦^C.

**Figure 2.**
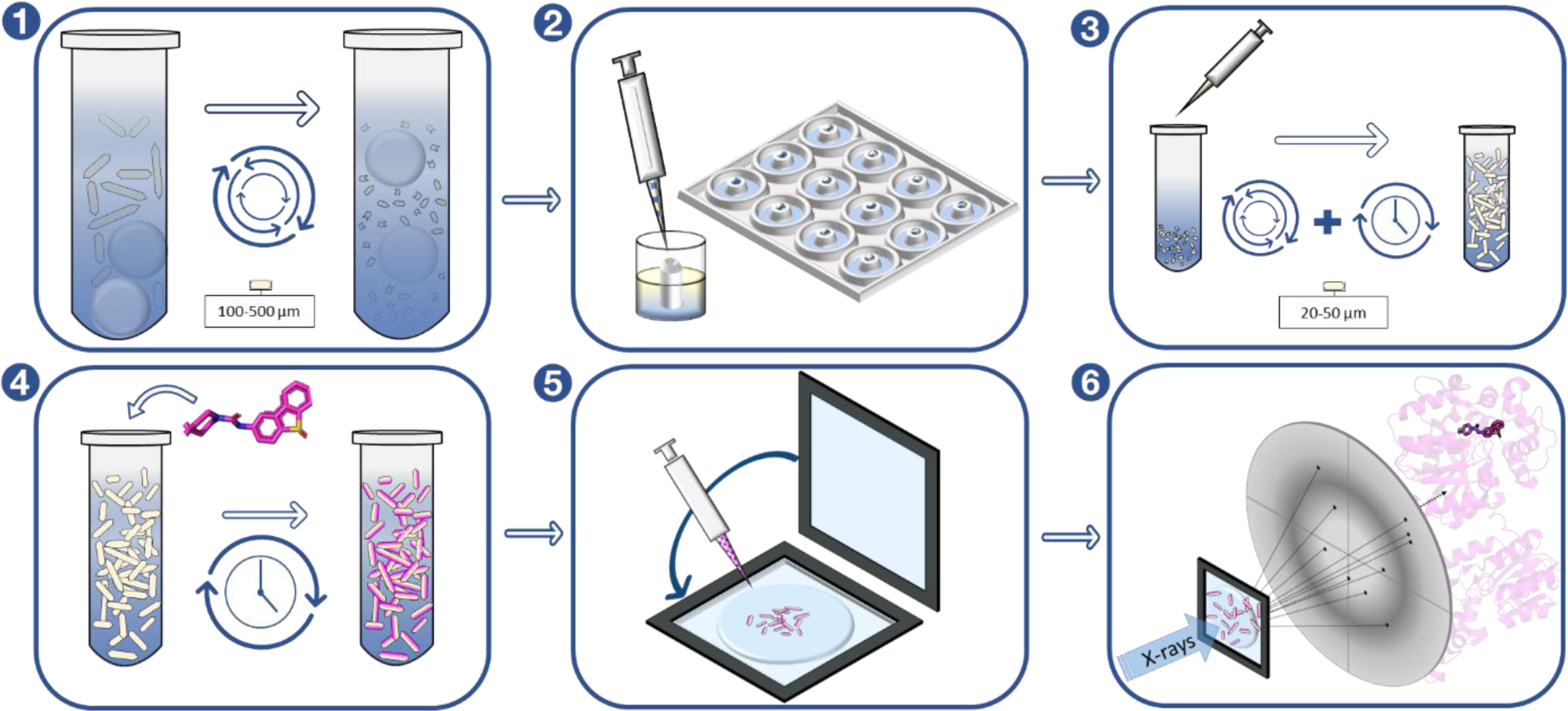
Microcrystallization and data collection workflow. **Panel 1:** Production of crystal seeds from macro crystals using seed beads and vortexing. **Panel 2:** Hybrid crystallization used for screening to find optimal batch crystallization conditions. **Panel 3:** Batch crystallization with seeding and vortexing to induce nucleation. **Panel 4:** Soaking of compound into the microcrystals. **Panel 5:** Dispensing of soaked crystals onto the chip and sealing. **Panel 6:** Fixed-target raster grid-scan data collection on ligand-soaked crystals at a synchrotron source.

Ligands dissolved in DMSO were added to a buffer (43.5 % (w/v) PEG 3350, 0.1 M Tris-HCl (pH 8.5) and 0.1 M Li_2_SO_4_), spun down to remove any precipitated ligand, and mixed into the crystal slurry to a final concentration of 10-50 mM, 2-72 hours before data collection.

### 2.3. Data collection and processing

All data collection was performed at the MX beamline BioMAX of MAX IV Laboratory using a fixed-target serial crystallography setup (Shilova *et al*., 2020). To increase the crystal hit-rate, the crystals were concentrated by centrifugation for 5-30 s at 1,500 x g. Next, 1.8-2 µl of the crystal solution were sandwiched between two Silson membranes (SiRN-5.0-200-2.5-1000) and manually mounted onto the goniometer using a reusable pin. Room-temperature diffraction data were collected on a Dectris Eiger 16M hybrid pixel detector by raster grid scanning using a 20×20 µm X-ray beam with 100 % transmission and an exposure time of 11 ms.

The datasets varied in size between 50,000 and 100,000 images. Hit finding, indexing and integration was done using indexamajig in the CrystFEL suite (White *et al*., 2012; White, 2019). Partialator was used for merging and scaling of the data. The resolution of each dataset was determined based on a CC_1/2_ of at least 30 and an I/σ(I) of at least 0.7. The structures were solved by molecular replacement in the ccp4i suit (Agirre *et al*., 2023) using a non-disclosed structure of sEH solved by rotational crystallography at cryo-temperature as a search model. The model was subsequently refined using Buster (Bricogne G. *et al*., 2017), while COOT (Emsley & Cowtan, 2004) was used for real-space rebuilding and refinement. Ligand restrains were generated with the programs writedict or Grade (Smart, O. S., *et al*., 2011; AFITT). The software PyMOL (Schrodinger, 2015) was used to produce figure of the structures. For data processing and refinement statistics see Table 1.

**Table 1:** Data collection, processing and refiner.

| Table 1<br>Data collection, processing and refiner |  |  |  |  |  |  |  |  |  |
| --- | --- | --- | --- | --- | --- | --- | --- | --- | --- |
|  | APO (8QVM) | Compound 1 (8QVL) | Compound 2 (8QVK) | Compound 3 (8QWH) | Compound 4 (8QWI) | Compound 5 APO (8QWG) | Compound 6 (8QVH) | Compound 7 (8QVG) | Compound 8 (8QVF) |
| *a*, *b*, *c* (Å) | 94.41 94.41, 247.57 | 94.39 94.39, 247.45 | 94.43 94.43, 246.89 | 94.38 94.38, 246.95 | 93.95 93.95, 246.69 | 94.4 94.4, 246.96 | 94.14 94.14, 247.26 | 93.87 93.87, 246.37 | 94.33 94.33, 246.92 |
| $\alpha$, $\beta$, $\gamma$ (°) | 90. 90. 120 | 90. 90. 120 | 90. 90. 120 | 90. 90. 120 | 90. 90. 120 | 90. 90. 120 | 90. 90. 120 | 90. 90. 120 | 90. 90. 120 |
| Total no. of reflections | 114534 | 122064109 | 3310882 | 10239340 | 77265832 | 1593253 | 8738291 | 16249553 | 977566 |
| No. of unique reflections | 45033 | 71073 | 72694 | 63407 | 69231 | 60947 | 61484 | 56223 | 63304 |
| Completeness (%) | 100 | 100 | 100 | 100 | 100 | 100 | 100 | 100 | 99.8 |
| Redundancy | 66.33 | 1717.446977 | 41.55 | 158.15 | 1116 | 26.14 | 93.32 | 289.019 | 15.442 |
| $\langle I/\sigma(I) \rangle$ | 3.66 (0.71) | 6.78 (0.79) | 5.96 (0.71) | 6.10 (0.72) | 9.42(1.62) | 5.47 (0.80) | 6.22 (0.70) | 6.20 (0.72) | 4.49 (0.83) |
| $CC_{1/2}$ | 95.13 (0.45) | 98.75 (42.49) | 94.24 (30.02) | 98.134 (52.85) | 99.29 (77.00) | 94.89 (32.86) | 98.86 (43.65) | 96.93 (48.33) | 97.07 (31.41) |
| *R*<sub>split</sub> (%) | 18.34 (96.91) | 10.07 (105.58) | 10.63 (144.13) | 11.48 (111.19) | 7.53 (50.98) | 16.88 (128.53) | 13.40 (132.62) | 12.80 (84.77) | 15.54 (151.65) |
| Overall *B* factor | 42.73 | 46.57 | 54.65 | 55.51 | 43.37 | 53.19 | 44.68 | 65.76 | 65.25 |
| Total no. of images | 84855 | 109977 | 41140 | 38108 | 94787 | 32446 | 43902 | 94426 | 98275 |
| Indexed images | 36503 | 72003 | 21078 | 18314 | 65236 | 6713 | 27369 | 37162 | 11305 |
| Structure refinement |  |  |  |  |  |  |  |  |  |
| Resolution range (Å) | 44.11 - 2.00 (2.00-2.01) | 38.81-2-14 (2.14-2.15) | 49.5-2.14 (2.14-2.15) | 49.26-2.20 (2.20-2.22) | 49.15-2.12 (2.12-2.14) | 44.09-2.20 (2.20-2.22) | 49.11 – 2.24 (2.24–2.25) | 38.6-2.20 (2.20-2.21) | 44.06-2.3 (2.30-2.32) |
| *R*<sub>work</sub>/*R*<sub>free</sub> | 20.25/24.64 | 19.15/22.56 | 19.27/23.43 | 19.05/22.54 | 17.92/20.64 | 20.59/24.08 | 19.07/22.29 | 21.92/25.39 | 20.81/26.67 |
| Waters | 187 | 131 | 133 | 104 | 181 | 114 | 139 | 182 | 61 |
| Bonds (Å) | 0.0081 | 0.00766 | 0.00787 | 0.00765 | 0.00843 | 0.00763 | 0.00755 | 0.00764 | 0.00752652 |
| Angles (°) | 0.9421 | 0.92591 | 0.93709 | 0.944 | 0.9351 | 0.93848 | 0.93325 | 0.95014 | 0.953738 |
| Average *B* factors (Å<sup>2</sup>) | 45.4 | 49.33 | 56.64 | 55.28 | 45.84 | 54.73 | 45.28 | 58.55 | 62.95 |
| Ramachandran plot |  |  |  |  |  |  |  |  |  |
| Favoured regions (%) | 96.76 | 97.37 | 97.36 | 97.39 | 96.5 | 96.65 | 95.93 | 97.02 | 95.54 |
| Outliers (%) | 0.38 | 0.19 | 0.38 | 0.37 | 0.39 | 0.74 | 0.37 | 0.74 | 0.56 |

## 3. Results and discussion

### 3.1. Crystallization & data collection

#### 3.1.1. Development of a new crystallization work-flow for SX

A key factor for a successful SX study is the ability to generate homogeneous microcrystals of high quality and in sufficient amounts (Dods *et al*., 2017). Thus, a robust crystallization protocol is required. For the purpose of generating the quantities needed, batch crystallization has typically been used for growing microcrystals (Beale *et al*., 2019). A severe limitation with using batch crystallization, however, is that it typically requires very large volumes of protein already at the screening stage. Another disadvantage is that visual inspection of crystallization progress is difficult. Our so-called hybrid crystallization approach, described below, presents a solution to both these problems.

Initial microcrystallization trials of sEH using vapor diffusion in sitting-drop plates with addition of seeds showed that reducing the protein concentration or increasing the concentrations of Li_2_SO_4_ and PEG3350 decreased the size of the crystals. In this study, we aimed for a crystal size of 20-40 µm in length as this was expected to give a good balance between high diffraction and low number of overlaps of crystals on the grids.

The main issue with the initial sitting-drop experiments was the low number of crystals generated in each drop. This is problematic as a low crystal density gives a low hit-rate, i.e., a low number of recorded diffraction images. It was possible to increase the number of crystals by several orders of magnitude by adding seed solution to the drop, however, this also induced the formation of precipitate. The problem was in part overcome by screening each new seed stock and testing dilutions between 1:16-1:256 of the seed stock in 43 % (w/v) PEG3350 0.1 M Tris-HCl (pH8.5) and Li_2_SO_4_, where 10 % (v/v) seed stock was added to the precipitant solution in each case, to ensure an optimal number of nucleation points. If available, a crystal counter would likely have improved the reproducibility of this step, as has been shown by others (Shoeman *et al*., 2023). Crystals could be detected after one hour at 20 ^◦^C. To reduce the speed of crystal formation and thereby potentially improve the crystal packing, crystallization experiments were also performed at 12 ^◦^C and 4 ^◦^C. This reduced the speed of crystal growth but also induced the formation of aggregates, thus all further set-ups were done at 20 ^◦^C.

Batch crystallization is typically the preferred method to produce large amounts of microcrystals. However, in the case of sEH, optimal crystallization conditions for vapor diffusion experiments with drop sizes of 1-3 µl did not translate well into batch conditions with final volumes of 50-100 µl. Since it is not feasible to screen a large variety of conditions in batch due to the high protein consumption, we developed an alternative method that we refer to as hybrid crystallization.

Hybrid crystallization is done in regular sitting-drop plates where the solution in the reservoir well is a mixture of the precipitant solution and the protein buffer in the same ratio as that used between the precipitant solution and protein in the drop. Therefore, the drop size will not change over the course of the experiment as in a regular vapor-diffusion setup. Instead, the experiment is very similar to a batch setup, but without risking that the small-volume drop dries out. The main advantage of this micro-batch setup is that sample consumption can be kept very low while screening for optimal crystallization conditions. Moreover, the hybrid crystallization approach allows the screening to be performed in plates which facilitates visual inspection during the course of the experiment. Finally, the hybrid crystallization method can take advantage of automated dispensing to make it less labor intensive. Using a drop volume of 4-10 µl and a reservoir volume of 500 µl, a large number of conditions were tested. It was found that the protein to precipitant/seed solution ratio influenced the thickness of the crystals (Figure 3) and thereby the resolution obtained. If the width of a single crystal was below 10 µm then the resolution in most cases would be 3 Å or worse. The optimal conditions identified were a precipitant solution of 34 % PEG3350, 0.1 M Tris-HCl (pH 8.5) and 0.1 M Li_2_SO_4_ with the addition of 10 % (v/v) seeds in combination with a protein to precipitant/seed solution ratio of 1:4. This condition could be successfully translated to larger-scale batch set-ups of 50-100 µl in tubes for microcrystallization of sEH. Microcrystals of sEH were sensitive to transportation, which resulted in crystals either melting or aggregating in transit. This was possibly due to temperature variations during shipping, and the best diffraction was obtained from crystals produced on-site at the synchrotron facility. However, to make SX viable for drug discovery applications, where the optimal working model relies on crystals being produced in-house and then sent to a synchrotron for remote data collection, it is crucial to develop a logistic chain that can maintain an environment where crystals can be shipped in solution or dispensed onto a chip.

**Figure 3.**
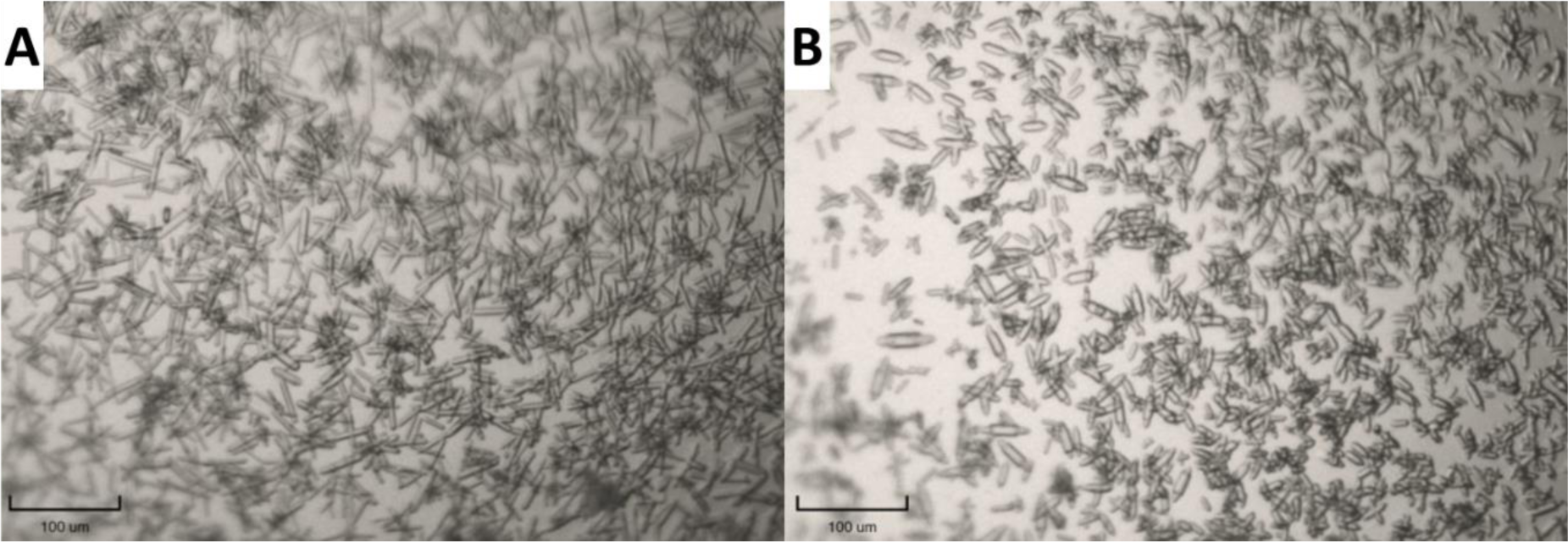
Microcrystals of sEH. The effect of using different protein:precipitant/seed solution ratios during screening for optimal conditions using the hybrid crystallization method is shown. **(A)** A protein:precipitant/seed solution ratio of 1 is used. **(B)** A protein:precipitant/seed solution ratio of 2 is used. A ratio of 2 gives a lower number of crystals but a more homogeneous sample with fewer overlapping crystals.

#### 3.1.2. Ligand introduction

For this study, we selected a subset of ligands that had previously been structural characterized in complex with sEH using conventional cryo-crystallography (Öster et al) in order to be able to do a direct comparison of the resulting structures. The selected ligands range from small weakly-binding compounds i.e., fragments, to larger potent inhibitors of sEH (Table S2).

Crystals of ligand-bound protein were prepared by soaking the crystals in high concentration (10-50 mM) of compounds dissolved in 43.5 % (w/v) PEG3350, 0.1 M Tris-HCl (pH 8.5) and 0.1 M Li_2_SO_4_ for a duration of 2 h or longer. We noted that several of the ligands were not fully solubilized under these conditions. While it is quite common in soaking experiments that the ligand precipitates, it is seldom detrimental in conventional cryo-crystallography where the crystal is fished out of the drop prior to data collection. However, in fixed-target SX experiments the crystal solution is added to a chip and thus ligand precipitate may result in high background and powder diffraction which affects the data quality negatively. Thus, more care must be taken when working with poorly soluble ligands in SX experiments. To reduce the problem caused by ligand precipitation, the ligand solution was centrifuged before addition to the crystal slurry.

#### 3.1.3. X-ray diffraction data collection

A number of different fixed-target chips were explored for the SX data collection where the best results were achieved using the Silson membranes. We found that an X-ray flux of at least 1×10^12^ photons/s with 100 % transmission were required for good diffraction. Each chip gave rise to 15,000-20,000 images on average and required 15-20 min to assemble and collect data from. Depending on hit-rate and the number of diffraction images aimed at, a complete dataset was compiled out of data collected from four to six chips. Thus, each dataset was acquired in about 1-2 hours. The high redundancy of the resulting data (up to 1700, see Table 1) suggests that the number of images could be drastically reduced, in particular in combination with a high crystal density as seen for e.g., crystals with compounds 1 and 2. Combined optimization of the crystal density and data collection strategy could thus significantly reduce the data collection time and thereby make SX a more feasible option from a synchrotron usage perspective. Room-temperature diffraction data were collected at the BioMAX beamline of MAX IV from crystals soaked with eight different compounds. Data collection, processing and refinement statistics are presented in Table 1.

### 3.2. Room-temperature structures of sEH and comparison with cryo-temperature structures

In this study, we present eight structures solved by SX at resolutions ranging from 2.1 to 2.5 Å resulting from sEH crystals soaked with different ligands. Complexes with the same ligands solved at cryo-temperature were reported to be of similar resolutions (1.9 – 2.5 Å). The microcrystals belong to the space group P6_5_22 which is the same as previously published sEH data. Unsurprisingly, the structures are overall very similar to structures solved using conventional cryo-crystallography.

#### 3.2.1. Ligand interactions

All the ligands with the exception of compound 5 could be clearly identified in the active site binding pocket (Figure 4) where they occupy the active site cavity similar to a large number of cryo-complex structures solved (Gomez *et al*., 2006; Öster *et al*., 2015; Shen & Hammock, 2012; Vázquez *et al*., 2023). Among the ligands, three compounds have a molecular weight below 300 Da and can be classified as fragments. For compound 1, two copies of the fragments could be identified in the active site (Figure 4). To find several copies of the same fragment bound in a structure is rather common in fragment screening due to the high compound concentrations used. Upon comparing the binding pose of each ligand in the SX structure with the corresponding cryo-structure, we conclude that the binding poses are very similar in all cases except one.

**Figure 4.**
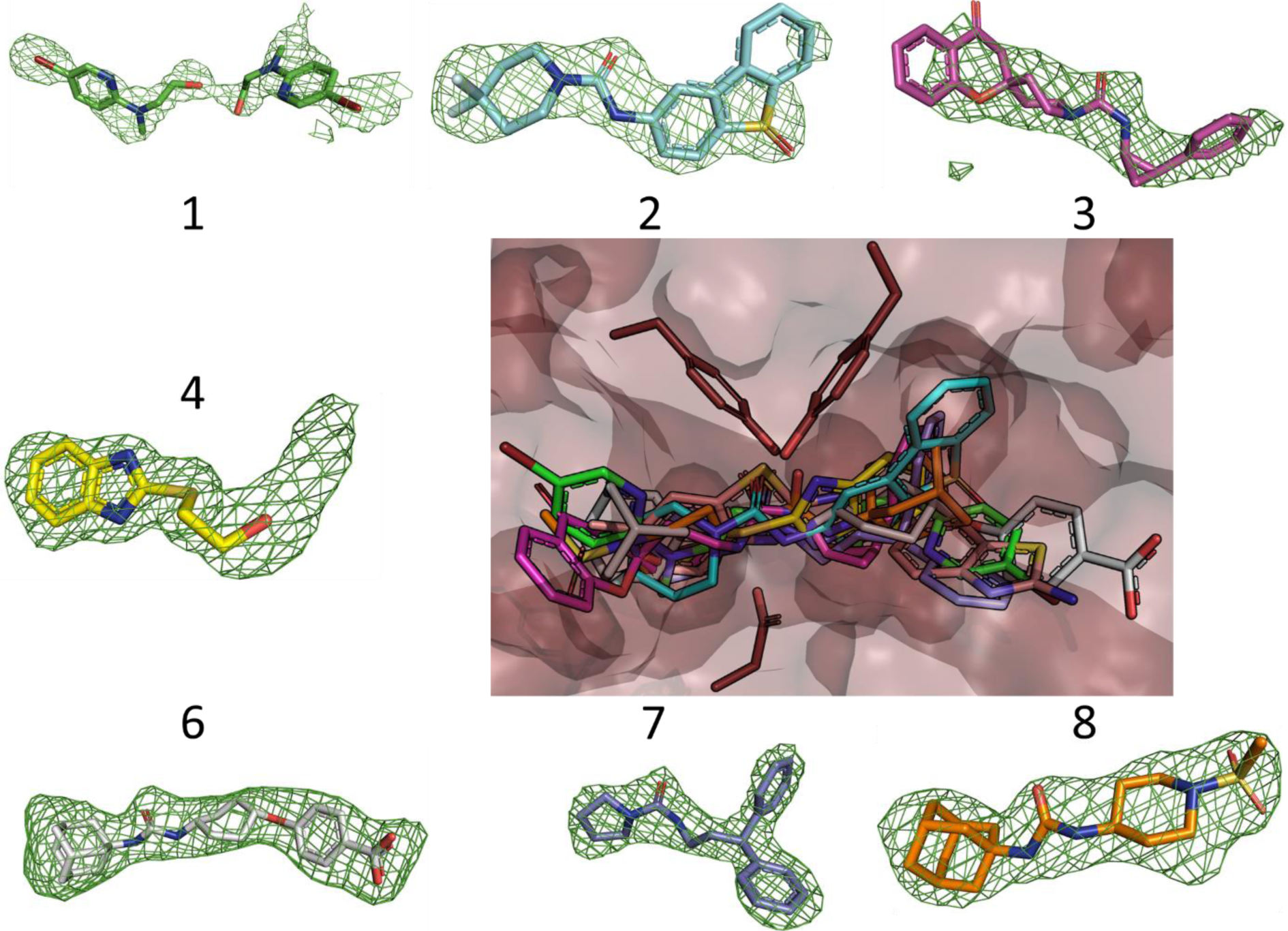
Protein-compound complex structures of sEH. The outer images show the different compounds that form complexes in their respective conformations numbered according to Table S2. The FoFc omit difference electron density map (green) is contoured at +3.5 σ in each case. The central panel displays an overlay of the seven room-temperature sEH complex-structures zoomed-in on the active site.

The structure generated from microcrystals soaked with compound 5 shows a large positive difference electron density feature in the active site. However, the compound cannot be reasonably modelled, and the density is instead well fitted with a PEG molecule (Figure 5A). The binding of a PEG molecule in the active site is also observed in the same position in the apo structure solved using SX at room temperature (Figure 5B). Interestingly, the room-temperature structure solved from crystals soaked with compound 5 is thus distinctly different from that solved at cryo-temperature, where compound 5 is well-fitted in the electron density at a position adjacent to the PEG position and not coordinated with the catalytic triad (Figure 5C). Moreover, in the cryo-structure, a sulphate ion and a water molecule reside where the density of the PEG molecule is found at room-temperature. To rule out the possibility that compound 5 binds at lower occupancy in the room-temperature structure, a model with the PEG molecule at 50 % occupancy and compound 5 in the adjacent position at 50 % occupancy was tested but results in a very poor fit of the electron density (Figure S1). We were also able to exclude the alternative that the relatively low number of indexed images for the compound 5 dataset (∼6,700, see Table 1) led to a poorly resolved active site ligand based on the fact that in an electron density map calculated using a sub-set of ∼5,000 of the compound 4 indexed images the compound 4 ligand is clearly visible. The fact that differences in ligand-binding features can be observed upon comparing structures solved at cryo- and room-temperature has been discussed previously (Fischer *et al*., 2015).

**Figure 5.**
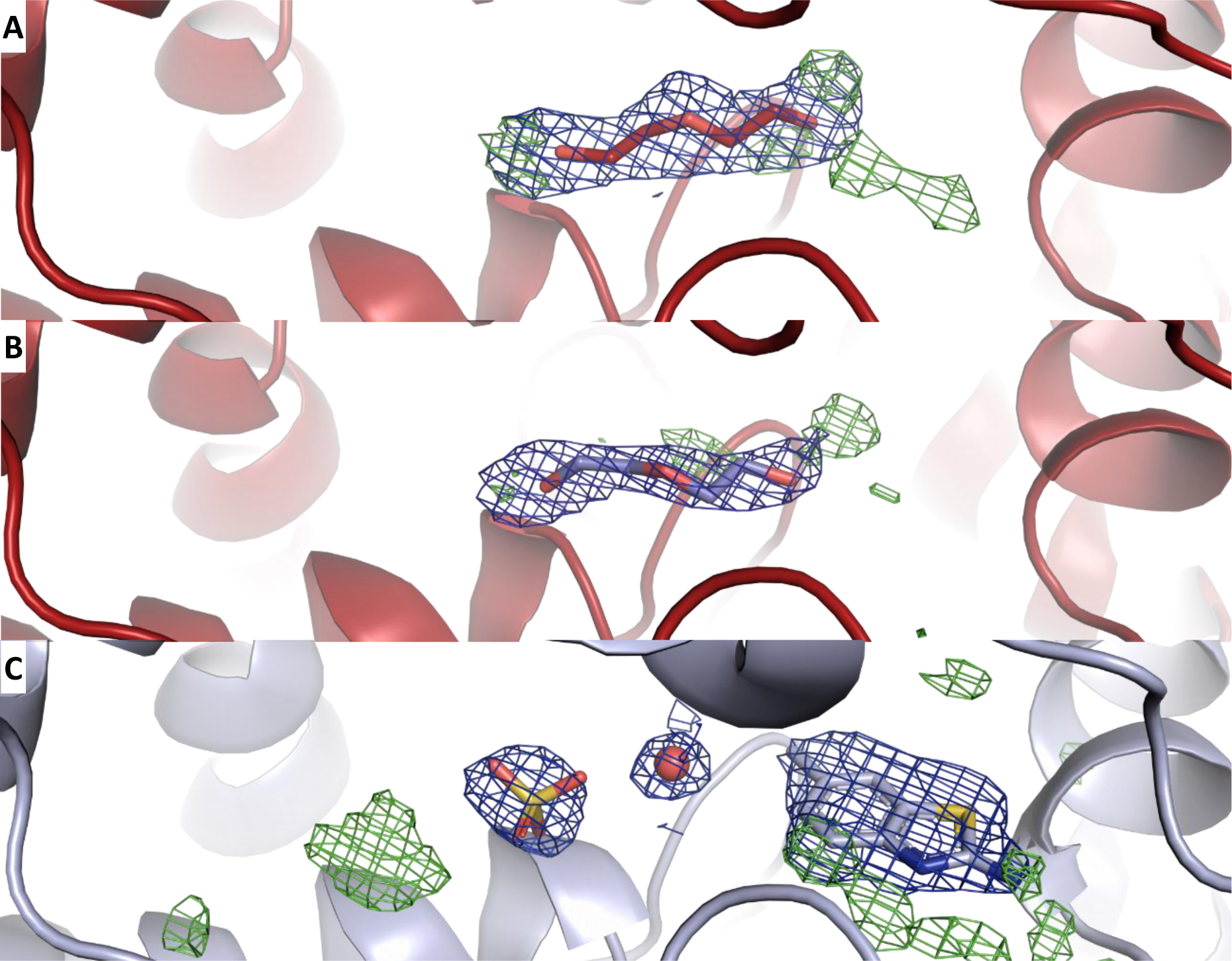
Temperature-dependent difference in fragment binding-mode. The room-temperature structure from crystals soaked with compound 5 is compared with the room-temperature apo structure and the cryo-structure in complex with compound 5. **(A)** In the room-temperature structure of sEH obtained from crystals soaked with compound 5 the active-site density is best fitted with a PEG molecule (PDB ID 8QWG). **(B)** The room-temperature apo structure is displayed with a PEG molecule bound in the active site (PDB ID 8QVM). **(C)** In the cryo-structure from crystals soaked with compound 5 (PDB ID 5AI8), the fragment is bound as well as a sulphate ion and a water molecule. For each structure, the 2FoFc electron density map is contoured at 1 σ (blue) and the FoFc electron density

#### 3.2.2. Water molecules, B-factors and loops

A significant difference between the sEH structures solved at room temperature and at cryo-temperature is the number of water molecules that can be modelled. The water content is three to five times higher in the structures solved using conventional cryo-crystallography (Figure 6). This is likely due to the fact that the elevated temperature increases disorder of water molecules that are not highly coordinated, as has been observed by others (Nakasako, 1999, 2001). Despite this difference in water content, we find that many well-ordered water molecules in the interior of the catalytic domain are conserved also in the SX structures.

**Figure 6.**
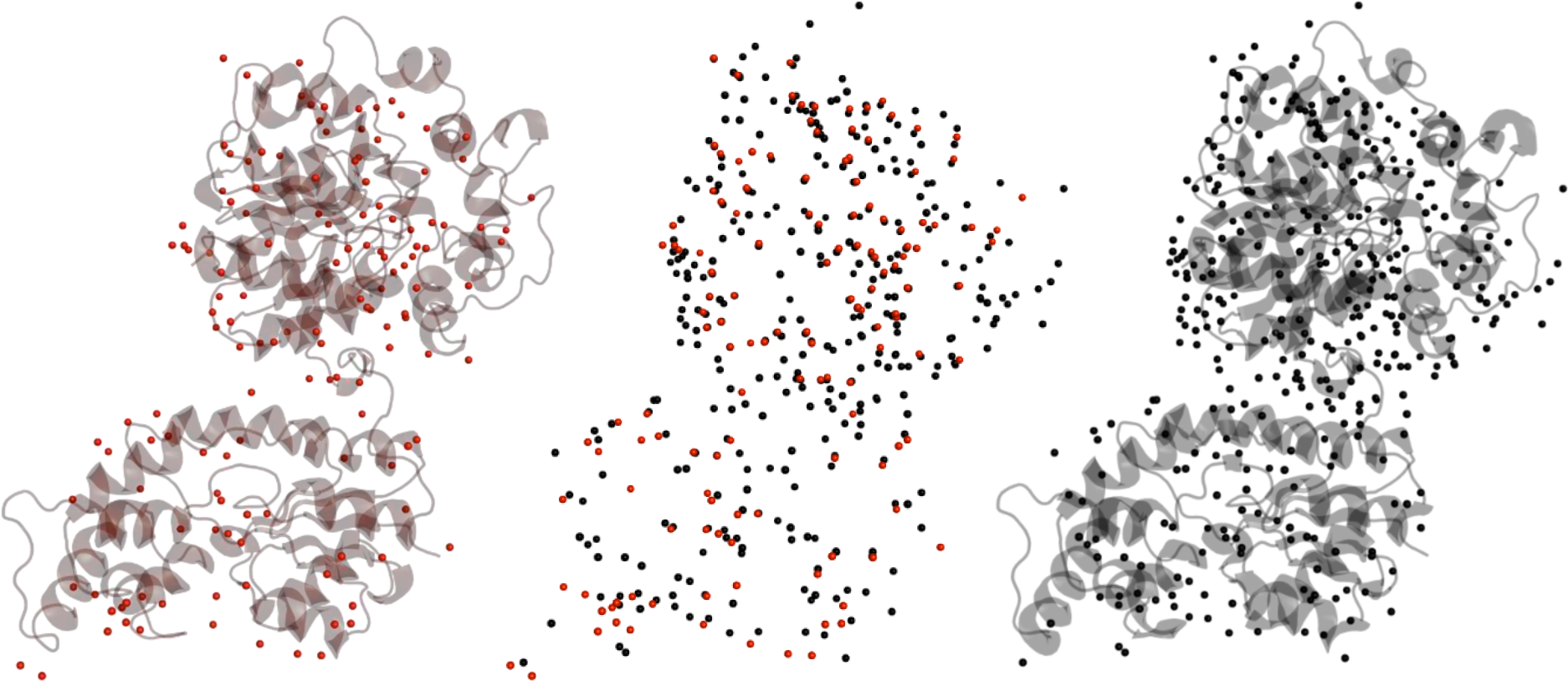
Water molecules modelled at room-temperature compared to at cryo-temperature. The left-hand side panel shows the room-temperature sEH structure (in complex with compound 2, PDB ID 8QVK) with water molecules in red. The right-hand side panel shows the corresponding cryo-structure (PDB ID 5AKE) with water molecules in black. The central panel displays an overlay of the water molecules.

As expected, we observe higher B-factors in the room temperature structures as compared to the cryo-temperature structures. This difference can be attributed to several factors, including increased thermal motion and enhanced flexibility when the structure is not cryo-trapped. The crystal lattice of microcrystals may be more resilient to conformational changes thus allowing the dynamics of the protein to be observed. Radiation damage can also be manifested in the form of high B-factors, and care must be taken to stay below the dose limit of 5 × 10^5^ Gy when X-ray radiation induced damage is typically observed in data collected at room temperature (Schneps *et al*., 2022). In our experiments, we calculate the radiation dose to ∼4.5 × 10^4^ Gy at each grid position of the chip (Bury *et al*., 2018). In a recent study of a cryptochrome using fixed-target SX, it was noted that the refined B-factors correlated well with those predicted from molecular-dynamics simulations (Schneps *et al*., 2022), pointing towards the protein being in a fully solvated, functional state in the microcrystals. The structures of sEH display flexibility in some loop-regions at both temperatures, including those surrounding the active site. Interestingly, the B-factors of the residues around the active site are significantly elevated in the room-temperature structures relative to the cryo-structures, suggesting a dynamic behaviour of the active site under ambient conditions that is masked in the frozen state (Figure 7). This type of enhanced loop mobility of functionally important regions was reported recently in a lower-resolution SX study (Schneps *et al*., 2022), where it was suggested that information gained from analysing dynamic behaviour could be useful to identify key structural elements in a protein.

**Figure 7.**
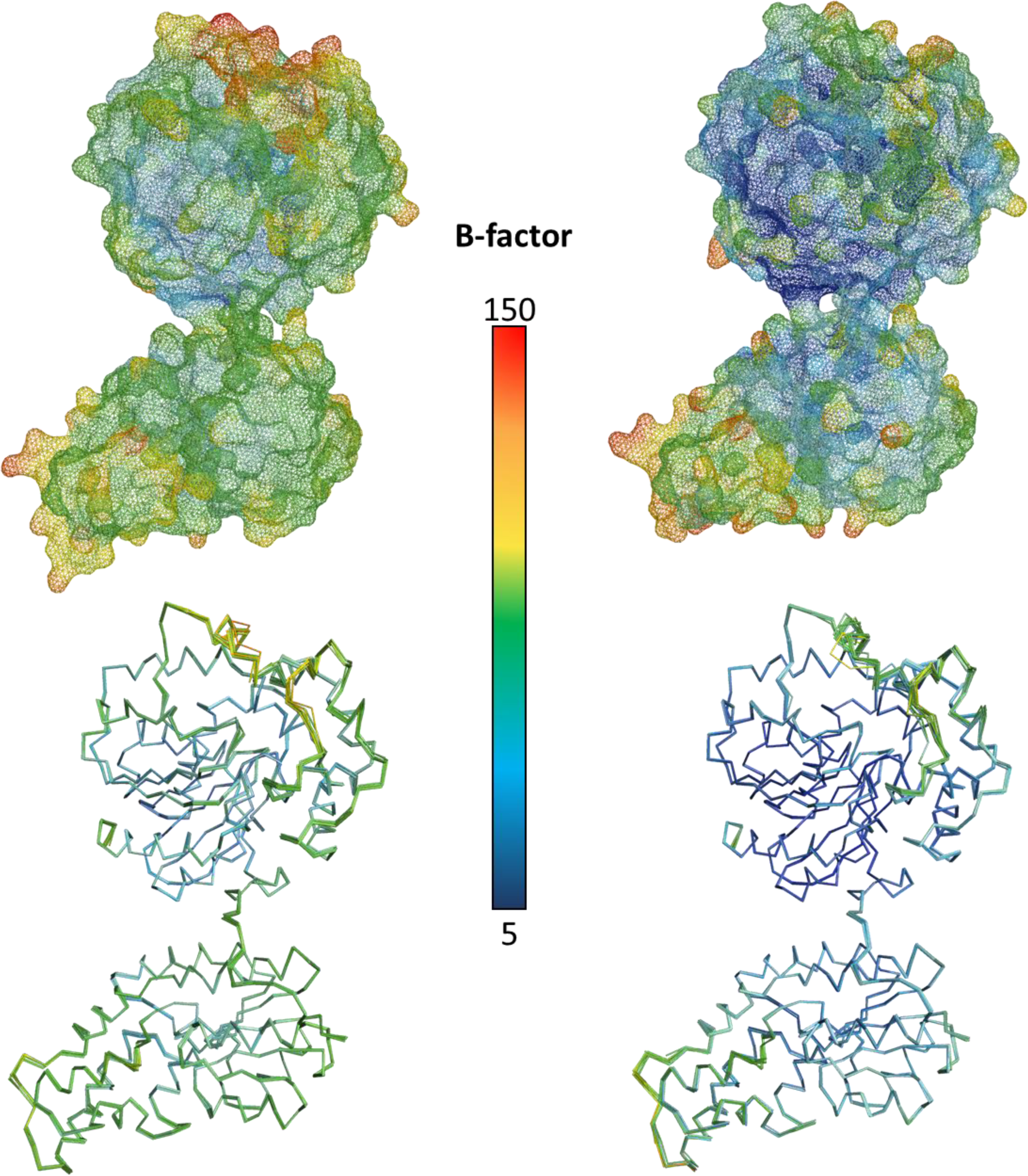
B-factor comparison. The B-factors of room-temperature (left-hand side) and cryo-temperature (right-hand side) sEH structures are plotted according to the central bar. At the top, a comparison is shown for one room-temperature structure (left, PDB ID 8QWG) and its cryo-temperature counterpart (right, PDB ID 5AI8). At the bottom, a backbone comparison of all the complex structures is displayed (room-temperature structures to the left, cryo-temperature structures to the right).

One of the most pronounced differences related to temperature in the structures of sEH is that several of the loops are more well-defined in the room-temperature models. Thus, it seems that it is possible to obtain more information on structural regions that are typically less ordered by use of SX. In our SX structures, we can confidently model the loop composed of residues 65 to 95 in the N-terminal domain, whereas this loop is either not possible to model at all in the cryo-structures or only associated with partial electron density (Figure 8). Interestingly, this loop is close to the putative phosphatase region of sEH (Cronin *et al*., 2003; Matsumoto *et al*., 2019; Vázquez *et al*., 2023). The fact that room-temperature diffraction data can provide superior electron density maps of flexible regions has been noted previously in a study of bacterial reaction centre using XFEL radiation (Dods *et al*., 2017), although the reason for this temperature effect is not clear. In contrast, it was recently shown that lipid tails and detergent molecules were not as well-resolved at higher temperature (Båth *et al*., 2022).

**Figure 8.**
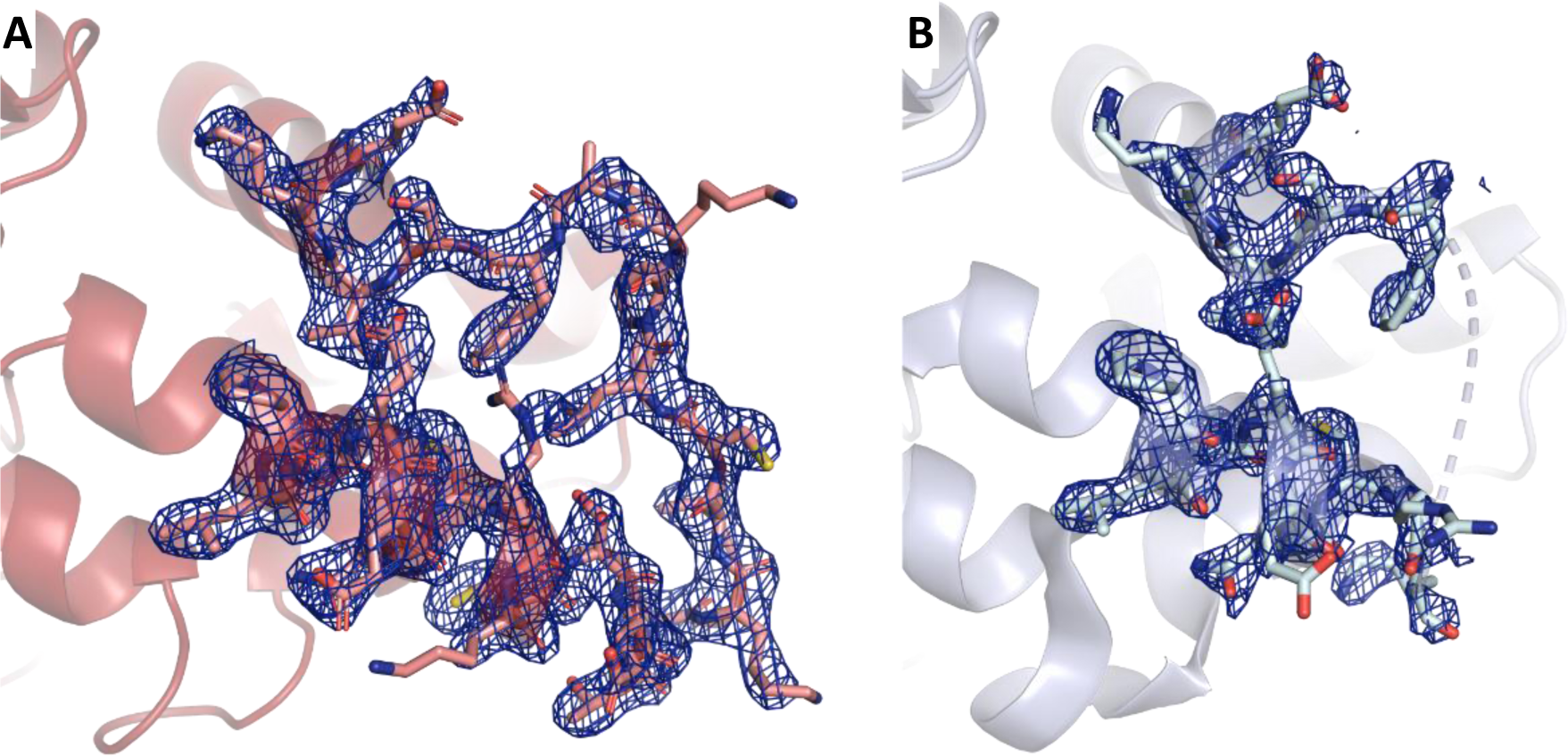
Ordering of loops. An example of a loop (Pro65 to Ala95) with higher degree of order in the room-temperature structures compared to in the cryo-structures. **(A)** The 2FoFc electron density map associated with crystals soaked with compound 5 at room-temperature is displayed (PDB ID 8QWG). **(B)** The 2FoFc electron density map associated with the corresponding cryo-structure is shown (PDB ID 5AI8). The 2FoFc electron density maps are contoured at 1 σ (blue).

## Conclusion

We have developed a robust workflow for efficient production of microcrystals for SX experiments. This involves the use of a hybrid crystallization approach to screen for suitable conditions using low amounts of protein. The method allows easy evaluation of crystallization trials and is readily scalable to larger volumes. Hybrid crystallization was successfully used to find microcrystallization conditions for the pharmaceutical target sEH, and based on this, larger amounts of microcrystals were produced by batch crystallization. We were able to solve room-temperature high-quality structures of sEH in complex with seven different ligands by serial synchrotron crystallography using a fixed-target set up. In comparison with previously published cryo-structures of the same protein-ligand complexes, we find that the ligands generally bind in similar fashion in the room-temperature and the cryo-structures. However, we observe one example of a temperature-dependent difference in the binding pose of a fragment, information that may influence the chemical design in a drug discovery project. In addition, we observe differences in protein conformations and solvent molecules both in the vicinity of the active site and in regions more remote. Thus, it may be beneficial to consider structural information at ambient temperature in a structure-based drug design project, in particular when working with weakly binding compounds.

Future studies will need to address challenges in working with poorly soluble ligands that cause precipitation, as well as issues with transporting and shipping crystals at room-temperature. Clearly, the most pressing challenge for SX from the perspective of pharmaceutical drug discovery is to tackle the slow turn-over and labor-intense nature of this method. Indeed, there is potential to optimize the SX workflow by developing a more automated crystal handling process. If successful, serial crystallography could become an attractive alternative also for high-throughput drug discovery campaigns.

## Acknowledgements

Crystal screening and X-ray diffraction data collection were carried out at the BioMAX beamline of MAX IV Laboratory (proposal numbers 20220186, 20210526, 20210912, 20200400, 20190933 and 20190465), at the P11 and P14 beamlines of PETRA III (proposal numbers 20190989 and MX-885, respectively), the PXI–X06SA beamline of Swiss Light Source (proposal number 20191098) and at the I24 beamline of Diamond Light Source (proposal numbers MX260673, MX29121 and MX24337). We are grateful for assistance by the beamline staff at the synchrotron facilities. We would like to thank Margareta Ek, Anna Aagaard, Lisa Wissler and Linda Öster, AstraZeneca, for valuable discussion regarding crystallization.

## Funding

GB acknowledges funding from the Swedish Foundation for Strategic Research (grant ID17-0060) and the Swedish Research Council (grants No. 2017-06734, 2021-05662 and 2021-05981).

## Author contributions

GB and HK conceived the experiment. Protein production, purification and crystallization was performed by AD, CP and JG. X-ray diffraction data collection was performed by AD, OU, MB, GW, HK and GB. X-ray diffraction data processing and refinement was performed by AD, GW, HK and GB. The manuscript was prepared by AD, HK and GB with additional input from all authors.

## Competing interests

GB is a co-founder of the company Serial X. HK is an AstraZeneca full-time employee and may own stock or stock options.

## Data and materials availability

The atomic coordinates and structure factors file for the sEH structure are available under accession codes 8QVM, 8QVL, 8QVK, 8QWH, 8QWI, 8QWG, 8QVH, 8QVG and 8QVF in the Protein Data Bank (www.pdb.org).

