## Supplementary material for "Exploring serial crystallography for drug discovery"

A. Dunge *et al.*

**This PDF file includes:**

Table S1
Table S2
Fig. S1

**Table S1**

**Crystallization**

| | | | |
|---|---|---|---|
| **Method** | Vapor-diffusion | Hybrid-crystallization | Batch-crystallization |
| **Plate type** | 24-well sitting drop plate | 24-well sitting drop plate | 0.5 mL Eppendorf Tubes |
| **Temperature (°C)** | 20 | 20 | 20 |
| **Protein concentration (mg/mL)** | 16 | 10-20 | 14 |
| **Protein buffer** | 20 mM Tris-Cl, pH 8.0, 100 mM NaCl, 10% glycerol and 1 mM TCEP | 20 mM Tris-Cl, pH 8.0, 100 mM NaCl, 10% glycerol and 1 mM TCEP | 20 mM Tris-Cl, pH 8.0, 100 mM NaCl, 10% glycerol and 1 mM TCEP |
| **Precipitant solution** | 32-38% PEG 3350, 0.1 M LiSO4, 0.1 M Tris pH 8.5. | 16-43% PEG 3350, 0.05-0.5 M LiSO4, 0.05-0.5 M Tris pH 8.5. | 34% PEG 3350, 0.1 M LiSO4, 0.1 M Tris pH 8.5. |
| **Volume and ratio of drop (p:w)** | Total volume 5-10 µL, 1:1 | Total volume 10 µL, 1:2 - 1:6 | Total volume 50-100 µL, 1:4 |
| **Crystallization reservoir** | 500 µL | 500 µL | - |
| **Seeding** | yes, streak seed | yes, 5-15% seed | yes, 10% seed (diluted 1:32-1:64) |

**Table S2**

**Compounds**

| Compound name | SMILES | MW (Da) | IC50 (uM) | Solubility (uM) | Heavy atoms | LogD pH7.4 (calculated) |
|---|---|---|---|---|---|---|
| **Compound 1** | CN(CCO)c1ccc(Br)cn1 | 231.09 | 331.7 | 0.9 | 12 | 2.11 |
| **Compound 2** | FC1(F)CCN(CC1)C(=O)Nc2ccc3c(c2)c4ccccc4S3(=O)=O | 378.4 | 0.015 | 5 | 26 | 2.4 |
| **Compound 3** | O=C(N[C@H]1C[C@@H]1c2ccccc2)N3CCC4(CC3)CC(=O)c5ccccc5O4 | 376.45 | 0.003 | 41 | 28 | 3.46 |
| **Compound 4** | OCCSc1nc2ccccc2[nH]1 | 194.26 | <1.37 | >1000 | 13 | 1.29 |
| **Compound 5** | Cc1ccc2nc(N)sc2c1 | 164.23 | 125.6 | 436 | 11 | 2.42 |
| **Compound 6** | OC(=O)c1ccc(O[C@@H]2CC[C@H](CC2)NC(=O)N[C@@]34C[C@@H]5C[C@@H](C[C@@H](C5)C3)C4)cc1 | 412.53 | 0.005 | 390 | 30 | 1.96 |
| **Compound 7** | O=C(NCCC(c1ccccc1)c2ccccc2)N3CCCC3 | 308.42 | <0.086 | 97 | 23 | 3.55 |
| **Compound 8** | CS(=O)(=O)N1CCC(CC1)NC(=O)NC23CC4CC(CC(C4)C2)C3 | 355.5 | 0.006 | 64 | 24 | 2.3 |

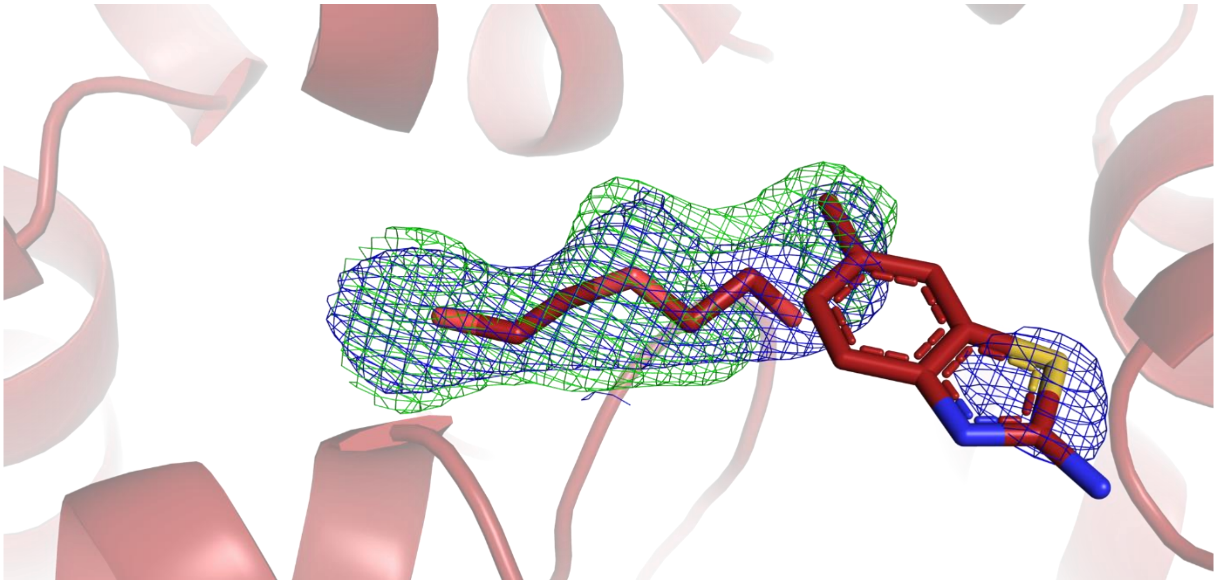

**Figure S1. The room-temperature structure of compound 5 soaked crystals.** The room-temperature structure of crystals soaked with compound 5 refined with 50 % occupancy of the PEG molecule and 50 % occupancy of compound 5. The resulting 2FoFc electron density map is contoured at 1 σ (blue) and the FoFc electron density map at +3.5 σ (green).
